# Thermodynamics Underpinning the Microbial Community-Level Nitrogen Networks

**DOI:** 10.1101/2024.05.21.595099

**Authors:** Mayumi Seto, Risa Sasaki, Hideshi Ooka, Ryuhei Nakamura

**Author notes:** **Corresponding author** Mayumi Seto, **Address:** Department of Chemistry, Biology, and Environmental Sciences, Nara Women’s University, Kita-Uoya Nishimachi, Nara 630-8506, Japan, **Email:**. **Author Contributions:** M.S., H.O., and R. N. conceived the work. M.S. and H.O. designed the theoretical formalism. M.S. developed the computational framework. M.S. and R.S. performed the numerical simulations and analyses. M. S., R.S, and H.O wrote the manuscript with input from R.N. **Competing Interest Statement:** Authors declare that they have no competing interests. **Data and code availability** All codes were written in the Wolfram Language platform using Mathematica 12. If the manuscript is accepted, all codes and data supporting the results will be archived in Dryad and the data DOI will be included at the end of the article.

## Abstract

Nitrogen species often serve as crucial electron donors or acceptors in microbial catabolism, enabling the synthesis of adenosine triphosphate (ATP). Although theoretically any nitrogen redox reactions could be an energy source, it remains unclear why specific reactions are predominantly utilized. This study evaluates energetically superior reactions from 988 theoretically plausible combinations involving 11 nitrogen species, oxygen gas, hydrogen ion, and water. Our analysis of the similarity between this model-based energetically superior network and the actual microbial community-level nitrogen network, reconstructed as a combination of enzymatic reactions, showed increased link overlap rates with thermodynamic weighting on reaction rates. In particular, existing microbial reactions involving solely nitrogen species and additionally oxygen, such as anaerobic ammonia oxidation (ANAMMOX) and complete and partial nitrification, were frequently identified as energetically superior among the examined reactions. The alignment of these reactions with thermodynamically favorable outcomes underscores the critical role of thermodynamics not only in individual metabolic processes but also in shaping the broader network interactions within ecosystems, consequently affecting biodiversity and ecological functions.

**Significance Statement:** This study advances our understanding of how thermodynamics governs energy metabolism at the community level within microbial ecosystems by systematically analyzing 988 potential redox reactions involving inorganic nitrogen species, oxygen gas, hydrogen ion, and water. We uncover that existing microbial reactions, such as anaerobic ammonia oxidation (ANAMMOX) and nitrification, stand out as energetically superior over other examined reactions. The robust alignment between model-predicted energetically favorable reactions and actual microbial nitrogen reactions underscores the predictive power of thermodynamic principles, even in ecological networks. Our findings extend the traditional applications of thermodynamics in biology, highlighting how thermodynamic constraints shape ecological networks and influence biodiversity and ecosystem functions in natural ecosystems.

## Introduction

Nitrogen forms compounds with oxidation states ranging from −3 to + 5 in the environment, serving as one of the key elements driving redox reactions on Earth (1). The majority of the nitrogen cycle processes are catalyzed by microbial enzymes, encompassing processes such as nitrogen fixation, nitrification, denitrification, anaerobic ammonia oxidation (ANAMMOX), and dissimilatory nitrate reduction to ammonium (DNRA) (2, 3). The nitrogen metabolism conducted by microbes is closely linked to ecosystem productivity, greenhouse effects, and the preservation of water quality. A comprehensive understanding of their functions is essential for predicting future changes in ecosystems and biochemical processes, as well as for formulating environmental conservation strategies (4–7).

Microbial nitrogen metabolism serves not only for the assimilation of nitrogenous macromolecules, such as proteins, but also facilitates energy acquisition through catabolic processes. There is mounting evidence that microbial metabolisms can utilize nitrogen species as electron acceptors or donors in diverse ways to synthesize ATP (8, 9). The Gibbs energy change of a reaction (Δ_*r*_*G*) plays a crucial role in predicting the directionality or feasibility of catabolic reactions (10, 11). This is exemplified by the prediction and subsequent discovery of processes like ANAMMOX and complete ammonia oxidation (COMMAMOX), which bypass traditional nitrification processes, and demonstrate the significance of thermodynamic calculations in understanding microbial nitrogen cycling (12–16). Nonetheless, the reasons why certain nitrogen reactions are favored as energy sources over a broad range of potential reactions remain elusive, and the full extent to which thermodynamics can predict microbial preferences for catabolic reactions at the community level has yet to be fully understood.

In this study, we systematically analyzed 988 reactions involving combinations of nitrogen half-reactions to identify those that are energetically advantageous and assess their alignment with existing microbial nitrogen reactions. Our approach comprehensively assess a wide range of nitrogen redox reactions, beyond a limited number of targeted reactions. Additionally, we provide a quantitative evaluation showing the role of thermodynamic properties in shaping the community-level nitrogen network using thermodynamic weighting on reaction rates and graph theoretical approaches. We simulated the inorganic nitrogen redox dynamics to identify a network of energetically superior reactions and compared it with microbial community-level nitrogen network, represented by the combination of microbial enzymatic functions. Notably, the reactions deemed energetically favorable by our model exhibited strong concordance with those integrated into the community-level network. These findings suggest that the microbial community-level nitrogen catabolic network has been shaped to align with increasing energy acquisition, significantly influenced by the thermodynamic properties of nitrogen species.

## Results

### Thermodynamic constraints on steady state nitrogen species composition

This model simulates the dynamics of 11 nitrogen species within an open, homogeneous single-box aquatic system (see Material and Methods for details). It incorporates 988 reactions derived from all combinations of 37 nitrogen species half-reactions and the reaction O_2_ + 4H^+^ + 4e^−^ ⇌2H_2_O, in addition to other reactions (see Supplementary Information and Data). Ammonia was selected as the sole reactive and reduced nitrogen source due to its abundant global flux from nitrogen fixation to the Earth’s biosphere (17) and its likely role as a predominant nitrogen source on early Earth from deep-sea hydrothermal vent emissions (18). The intentional exclusion of other redox-sensitive elements such as carbon and sulfur was designed to facilitate evaluations of extent to which the thermodynamic properties of nitrogen themselves can affect microbial preferences for catabolic nitrogen reactions.

The reaction rate *r*_*i*_ for each of the 988 reactions was modeled based on mass action kinetics. The Gibbs energy change of the reaction serves as not only an indicator of whether the reaction can be an energy source for ATP synthesis but also whether the reaction proceeds in the forward or backward direction; a reaction with a negative and lower Δ_*r*_*G* proceeds relatively quickly. To highlight this relative effect of Δ_*r*_*G* on reaction rates, we introduced a thermodynamic weighting coefficient (*p*) to directly adjust the influence of the change in Gibbs energy on reaction rates. As *p* increases, the difference in reaction rates between the forward reactions with Δ_*r*_*G* < 0 and their backward reactions becomes more pronounced. The model assumes a uniform effect of activation energy across all reaction coefficients (*k*_*i*_), represented by the reference reaction rate constant *k*_*def*_. This idealization enables us to specifically assess the significance of thermodynamic properties alone on the overall dynamics of nitrogen redox system and on the similarity between the energetically favorable reactions and microbial catabolic reactions.

The presence and magnitude of thermodynamic weighting significantly influenced the steady-state composition of nitrogen compounds by affecting the reactivity of nitrogen species (Fig. 1). The absence of thermodynamic weighting (*k*_*i*_ = *k*_*def*_) or low *p* values underestimates the rates of energetically favorable reactions that generally deplete thermodynamically reactive compounds. Consequently, reactive nitrogen species such as hydroxylammonium ions accumulated even in oxidative environments (Fig. 1a, b). However, as *p* increases, reactions with small Δ_*r*_*G* proceed relatively quickly, resulting in the prevalent nitrogen species under certain oxygen and pH levels reflecting those on present-day Earth, where thermodynamically stable nitrogen in the form of N_2_ is abundant (Fig. 1c, d and Fig. S1). An increase in O_2_ levels enhanced the accumulation of oxidized nitrogen forms like NO_3_^−^, NO_2_^−^, and N_2_, while reducing forms like NH_2_OH, NH_3_OH^+^, NH_3_, and NH_4_^+^ decreased. Rising hydrogen ion levels favored the formation of NH_3_OH^+^ and NH_4_^+^ due to protonation. The distribution patter of nitrogen species concentrations was largely consistent across various NH_3_ supply levels (Figs. S2).

**Figure 1.**
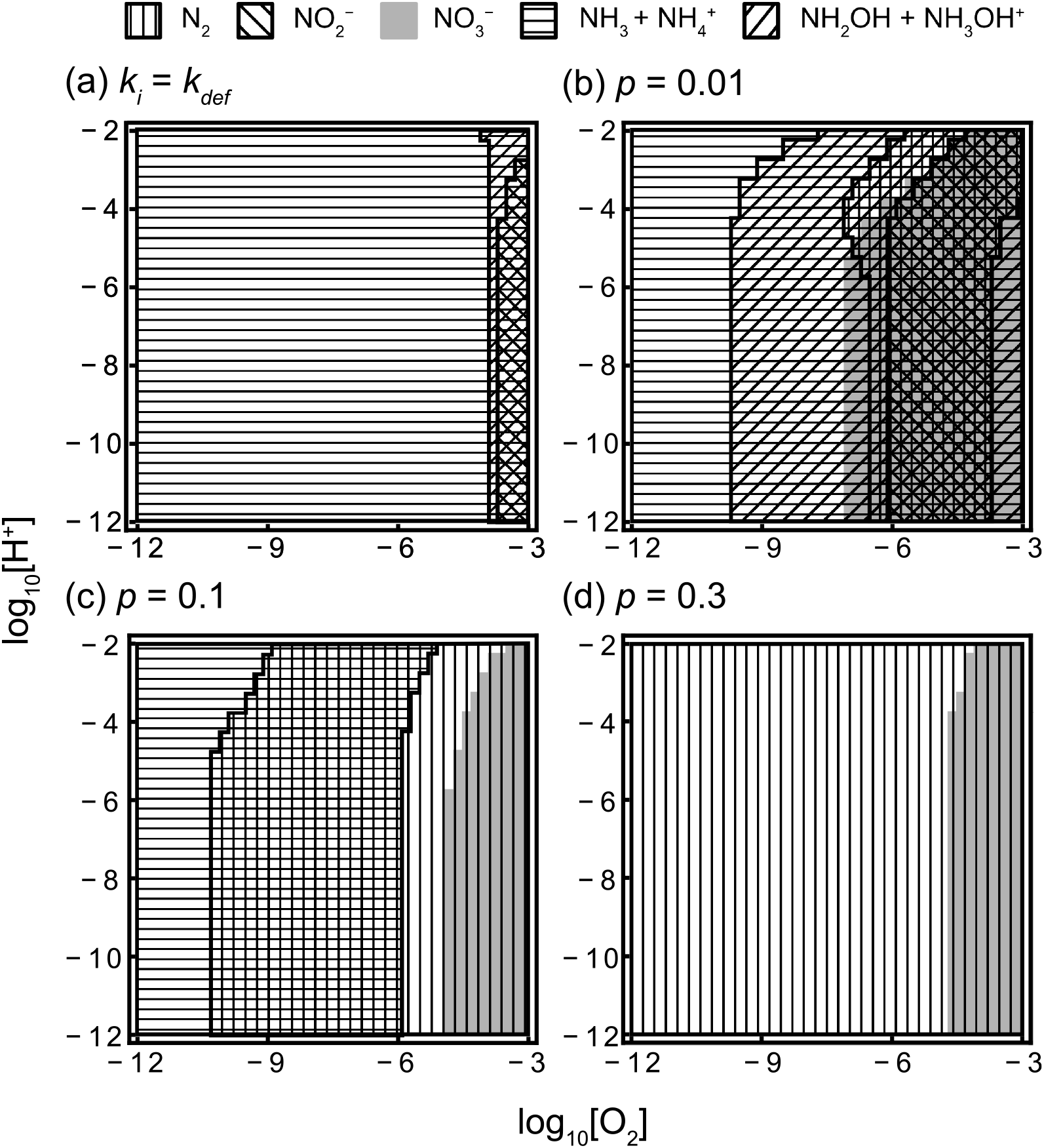
The combinations of major nitrogen species at each oxygen and hydrogen ion concentration in the steady state. The term “major nitrogen species” refers to those that constitute more than 1% of the total concentration of the 11 nitrogen species. Each nitrogen species exists as a major nitrogen species in the respective patterns shown in the legend. (a) represents the scenario without considering thermodynamic weighting on the reaction rates: 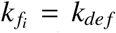. (b, c, d) correspond respectively to when *p* = 0.01, 0.1, and 0.3. *I* = 10^−5^, *D* = 10^−3^, *T* = 288, and *k*_*def*_ = 1.

### Model-derived and microbial community-level nitrogen networks

We assessed the similarity between model-derived energetically superior reactions and actual microbial catabolic reactions by representing the nitrogen reactions as chemical networks using a graph-based approach. We defined the graph *G*(*V, E*), which consists of 11 nitrogen species as nodes *V* and the directed links *E*, representing the nitrogen species transformation through 988 reactions (Figure 2a). The directed link set *E*_*i*_ for each reaction *rxn*_*I*_ includes all reactant-byproducts pairs. For instance, the reaction 1 NO_2_^−^ + 0.5 O_2_ → 1 NO_3_^−^ yields a single directed link {(NO_2_^−^, NO_3_^−^)}. The overall set *E* is the aggregation of directed links from all 988 reactions: 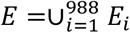. Each *rxn*_*i*_ is characterized by its steady-state Gibbs energy change Δ_*r*_*G*_*i*_, reaction rate *r*_*i*_, and power *P*_*i*_ = −Δ_*r*_*G*_*i*_ × *r*_*i*_, representing the energy output per unit time. By assigning one of the energy rating values to each *rxn*_*i*_ and selecting energetically superior reactions, we delineate a subset of nitrogen species nodes *V*_*ω*_ and directed links *E*_*ω*_, with the subscript *ω* representing *P, r*, or −Δ_*r*_*G* (Fig. 2b). Hereafter, *G*’(*V*_*ω*_, *E*_*ω*_) will be referred to as ‘the energetically superior network’.

**Figure 2.**
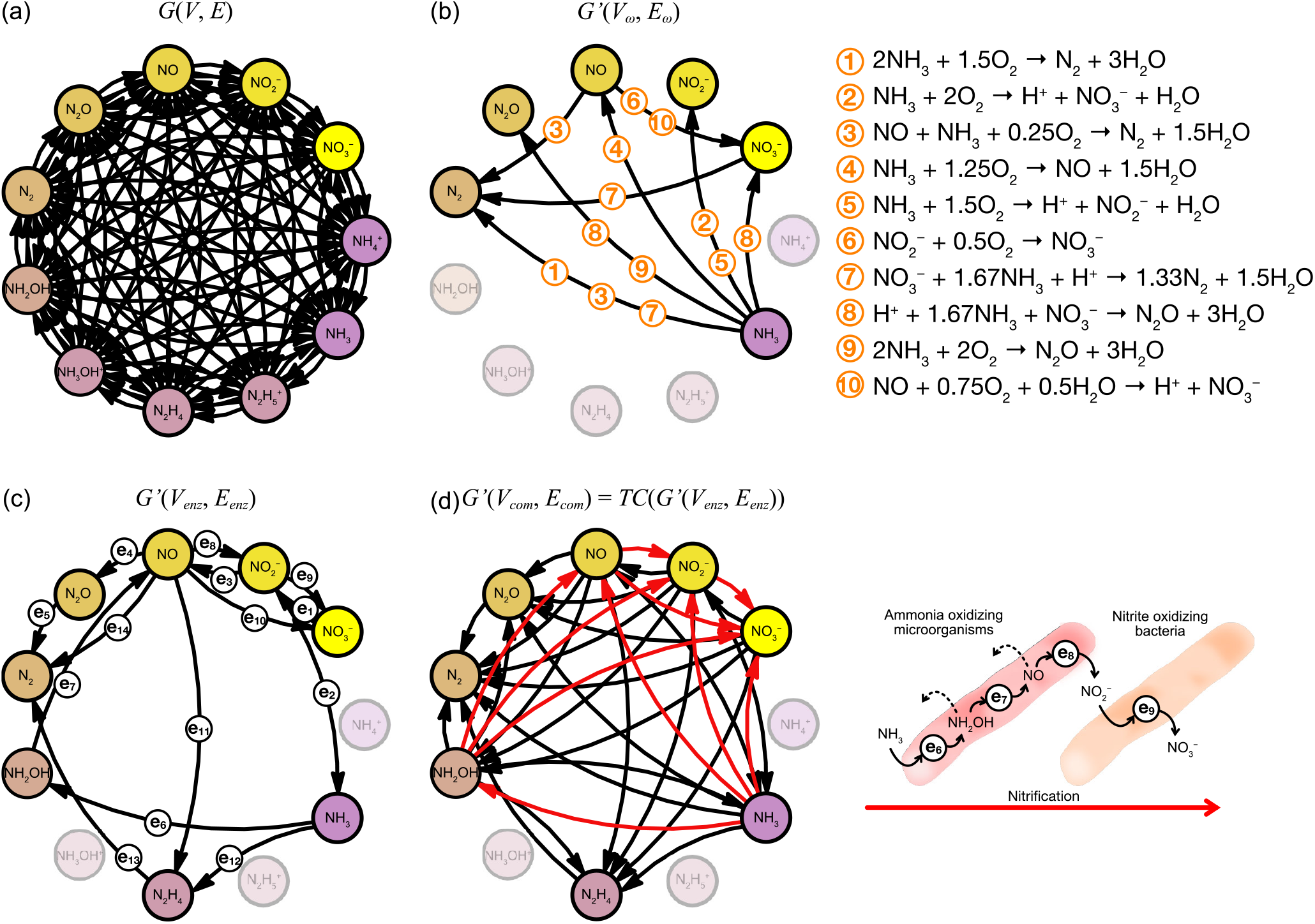
Nitrogen reactions as graphs. (a) The nitrogen reaction network comprising nitrogen species as nodes *V* with their transformations represented as directed links *E*, derived from the 988 reactions. (b) The energetically superior network formed from the top 10 reactions, ranked based on power *P*_*i*_, under steady state conditions with [O_2_] = 10^−5^ M, pH = 8, *p* = 0.3, *I* = 10^−5^, *D* = 10^−3^, *T* = 288, and *k*_def_ = 1. (c) The enzyme-level network, formed from enzymatic reactions as reviewed by Kuypers et al. (2018), excluding nitrogen fixation. Each link *e*_*i*_ corresponds to a nitrogen transformation link for a specific enzymatic reaction. (d) The community-level network, depicted as the transitive closure of the enzyme-level network. Red arrows show the transitive closure of the links from *e*_6_ to *e*_9_, which correspond to partial and complete nitrification.

The fundamental units of the microbial nitrogen reactions are enzymatic reactions, with microbes executing sequential or partial nitrogen transformations through the expression of specific enzymes. Microbial catabolic enzymatic reactions specifically targeting energy acquisition from nitrogen substances can also form a subgraph *G’*(*V*_*enz*_, *E*_*enz*_) (Fig. 2c). The directed links *E*_*enz*_ = {*e*_1_, …, *e*_13_} represent the nitrogen transformations by 13 of enzymatic reactions (e.g., *e*_1_ = (NO_3_^−^, NO_2_^−^)), as reviewed by Kuypers et al. (2018), excluding nitrogen fixation. The specific combinations of links e_1_ through e_13_ correspond respectively to DNRA (*e*_1_ and *e*_2_), denitrification (*e*_3_, *e*_4_, and *e*_5_), nitrification (*e*_6_, *e*_7_, *e*_8_, and *e*_9_), and ANAMMOX (*e*_3,_ *e*_11_, *e*_12_, and *e*_13_). The role of ammonium (NH_4_^+^) in nitrification is still unclear (19); however, the enzymatic reaction catalyzed by ammonia monooxygenase (e_6_ in Fig. 2c) is assumed to specifically convert NH_3_ to NH_2_OH, rather than NH_4_^+^ to NH_2_OH (20, 21). Hereafter, *G’*(*V*_*enz*_, *E*_*enz*_) will be referred to as ‘the enzyme-level network’.

Each microbial species possesses a subset of these enzymes. For example, ammonia-oxidizing microorganisms (AOM) oxidize NH_3_ to NO_2_^−^ using enzymes denoted by e_6_, e_7_, and e_8_ (22). The cooperation between AOM and nitrite-oxidizing bacteria (NOB), which oxidize NO_2_^−^ to NO_3_^−^ using e_9_, completes the nitrification process, while intermediate products of AOM, NH_2_OH and NO, may leak out, enabling additional transitions from NH_3_ to NH_2_OH and from NH_3_ to NO. Consequently, the microbial nitrogen network, formed through the integration of multiple enzymes within the community-level, should manifest as all possible combinations of enzymatic functions, represented by *G’*(*V*_*com*_, *E*_*com*_) (Fig. 2d). *G’*(*V*_*com*_, *E*_*com*_) is the transitive closure of *G’*(*V*_*enz*_, *E*_*enz*_) and identified using the TransitiveClosureGraph function in Mathematica 12. *G’*(*V*_*com*_, *E*_*com*_) will henceforth be referred to as ‘the community-level network’.

When the set of links *E*_*ω*_ in Fig. 2b matches the set *E*_*com*_ in Fig. 2d, each link within *E*_*ω*_ is associated with either an enzymatic reaction or their composite reactions.

Reaction 1 in Fig. 2b can be considered as a composite reaction of nitrification and ANAMMOX, or nitrification and denitrification. While the coexistence of these processes has been reported (23–26), recent evidence suggests the presence of microbial candidates that undergo these reactions at the individual level, termed as direct ammonia oxidation with a potential new enzyme (DIRAMMOX) (27–29). Reaction 2 is identified as complete nitrification, while 4, 5, 6, and 10 are its partial reactions. Reaction 3 resembles ANAMMOX but involves the utilization of oxygen, which can be considered as ANAMMOX occurring with partial nitrification of NO to NO_2_^−^ by using O_2_ as an electron acceptor. Reactions 7 and 8 can be considered ANAMMOX performed after reducing NO_3_^−^ to NO_2_^−^ using the reducing power of NH_3_. Reaction 8 and 9, with N_2_O as the final product, are associated with partial denitrification.

### The overlap rate between the model-derived and microbial community-level nitrogen networks

At each given O_2_ and pH conditions, we assessed the similarity between the energetically superior network and both the enzyme-level and community-level networks by comparing the edge overlaps:

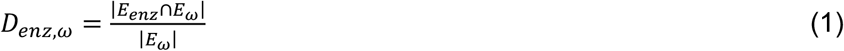

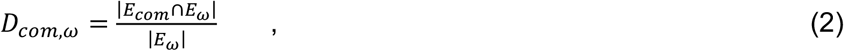

where |*E*| represents the number of links within *E*. For example, in Fig. 2b, the links derived from top 10 reactions ranked by power *P* (|*E*_*P*_| = 8) completely match the community-level links (|*E*_*com*_ ∩ *E*_*P*_| = 8), resulting in *D*_*com,P*_ = 1. A higher overlap rate suggests that the substructures within the enzyme-level or community-level network are supported by energetically superior reactions. To ensure consistency when comparing link overlaps across varying O_2_ and pH conditions, we initially selected an identical number of reactions, specifically the top 10, ranked by either *P, r*, or −Δ_*r*_*G*, to identify the set of links *E*_*ω*_.

Among three rating factors, under the same O_2_ and pH conditions, the overlap rates of links selected based on −Δ_*r*_*G* were lower than those based on *P* and *r* (Fig. 3(a)), suggesting that the existing microbial catabolic processes do not always prefer reactions with lower Δ_*r*_*G*. Reactions with lower Δ_*r*_*G* often involve thermodynamically reactive N_2_H_4_ and N_2_H_5_^+^ as reactants. However, maintaining these reactions at a steady state can be challenging because N_2_H_4_ and N_2_H_5_^+^ are quickly depleted, significantly reducing the reaction rates and power generation at steady state. Consequently, microbial catabolism may not sustainably rely solely on these reactions. The ANAMMOX process is currently the only microbial process that utilizes N_2_H_4_ as an intermediate (Fig. 2c), facilitated by electron recycling within the cell (30). This suggests that biological utilization of N_2_H_4_ or N_2_H_5_^+^ might fundamentally require elaborate mechanisms to maintain their concentrations.

**Figure 3.**
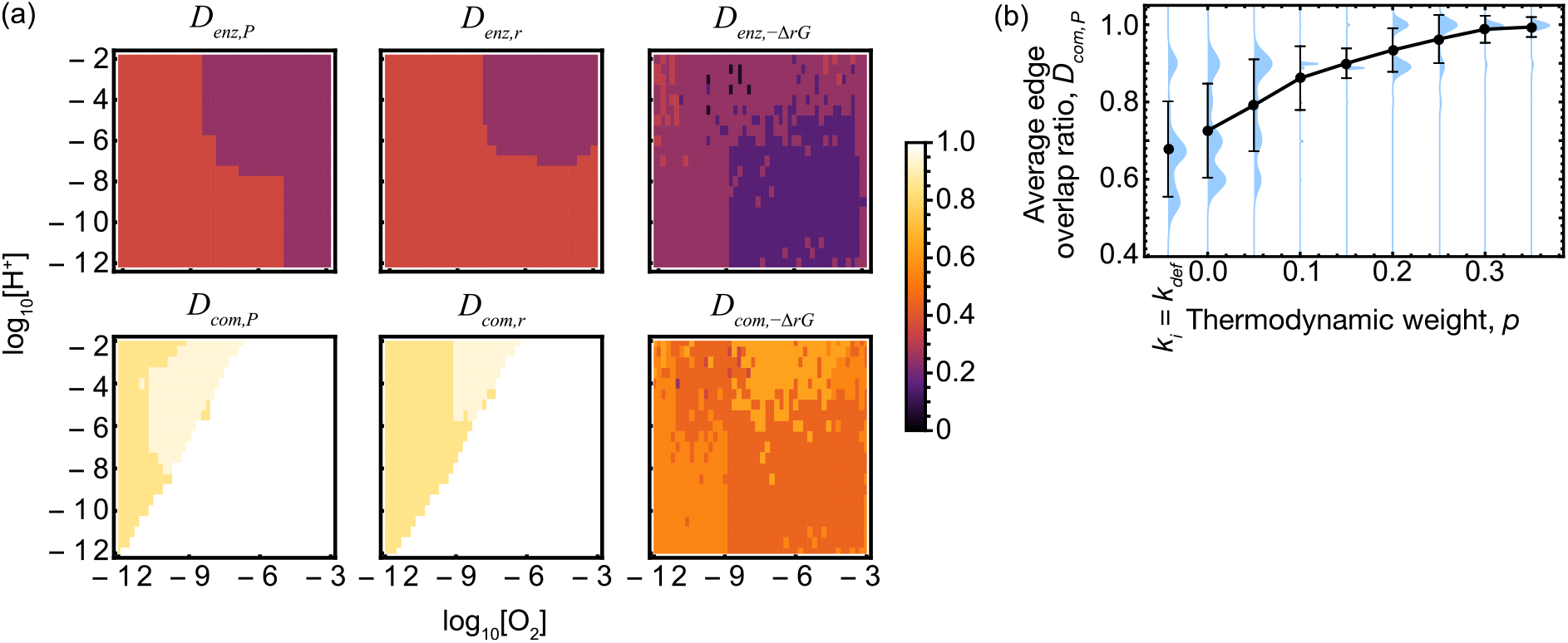
Edge overlap rates between the enzyme-level or community-level network and the energetically superior network, constructed from the top 10 reactions ranked by power (*P*), rate (*r*), and negative Gibbs energy of the reaction (−Δ_*r*_*G*). We denote these overlaps as *D*_*enz*,ω_ and *D*_*com*,ω_, where ω = *P, r*, or − Δ_*r*_*G*. (a) Edge overlap rates for each pH and oxygen concentration, with the top and bottom rows showing *D*_*enz*,ω_ and *D*_*com*,ω_, respectively. The columns from left to right correspond to energy rating factors by *P, r*, and −Δ_*r*_*G*, respectively. *p* = 0.2. (b) Average edge overlap rate *D*_*enz,P*_ in relation to the thermodynamic weight *p*, and in the absence of thermodynamic weighting (*k*_*i*_ = *k*_def_). The average edge overlap is calculated as the mean of edge overlaps under various oxygen concentrations and pH conditions. *I* = 10^−5^, *D* = 10^−3^, and *T* = 288. *k*_def_ = 10^−10^ for (a) and *k*_def_ = 10^−15^ for (b).

For all rating factors, *D*_*com,ω*_ consistently surpasses *D*_*enz,ω*_ across all O_2_ and pH conditions. This suggests that the nitrogen reactions within the community-level network are more likely to be characterized by energetically superior reactions than the enzymatic reactions that form its basis. Notably, the set of links *E*_*ω*_ at higher O_2_ levels significantly overlapped with the community-level links and were particularly involved in the complete or partial nitrification processes (cf. Fig. 2c). The reduced overlap in low O_2_ environments likely overlooks the role of other elements that provide reducing power, as DNRA and denitrification processes utilize organic matter, ferrous ion, hydrogen gas, hydrogen sulfide, and methane as electron donors (31–34).

Upon analyzing the response of the average *D*_*com,W*_ across all O_2_ and pH conditions to variations in the thermodynamic weight *p*, it was observed that the average *D*_*com,P*_ increased as *p* increased (Fig. 3b). In this model, the steady state of the system is predominantly influenced by the thermodynamic stability of nitrogen species at elevated *p* values. Although the behavior of nitrogen in the actual environment is influenced by more complex interactions with multi-element compounds, the increase in *D*_*com,P*_ supports the idea that microbial nitrogen catabolism and the structure of the resulting nitrogen community-level catabolic network are significantly influenced by the thermodynamic stability of nitrogen species themselves.

## Discussion

### Reactions with significant contributions to power generation

Since only a limited number of reactions contributing substantially to the system’s total power 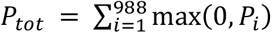 (Fig. S3), we identified key reactions from the top ranked reactions that collectively contribute to *θP*_*tot*_, where 0 ≤ *θ* ≤ 1. As *θ* approaches 1, the number of key reactions increases. However, the selection narrows as thermodynamic weight (*p*) increases, due to the dominance of reactions influenced by the thermodynamic stability of nitrogen species, even when *θ* = 0.9999 (Figure 4a).

**Figure 4.**
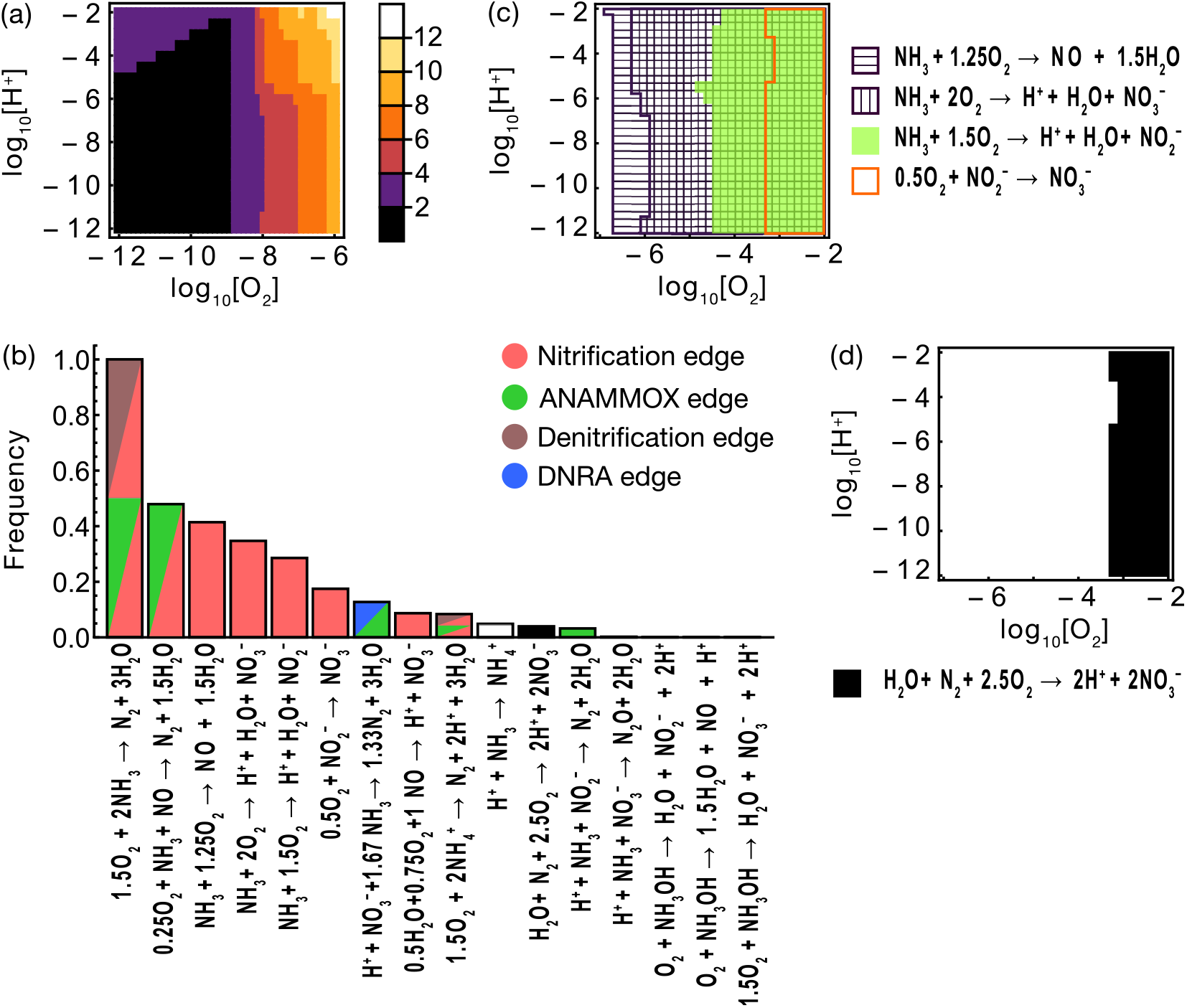
The set of reactions contributing to *θ* (0 ≤ *θ* ≤ 1) of the total power *P*_*tot*_. (a) The number of reactions *n* that represent 99.99% of the total power *P*_*tot*_ under 966 combinations of oxygen concentrations and pH conditions. (b) The frequency of occurrence of each reaction under 966 combinations of oxygen concentration and pH conditions. The color of the bars indicates the relationship between the reactions and the corresponding enzyme function links; white and black denote reactions not included in enzyme links. (c) Conditions of oxygen concentration and pH under which complete ammonia oxidation and its partial reactions significantly contribute to the total power. (d) Oxygen concentration and pH conditions where the oxidation of N_2_ to 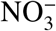 significantly contributes to the total power. *p* = 0.2, *k*_*def*_ = 1, *I* = 10^−5^, *D* = 10^−3^, *θ* = 0.9999 and *T* = 288K.

Fig. 4b highlights key reactions across 966 O_2_ and pH conditions, indicating the frequency with which each reaction was identified as energetically superior. A composite reaction of nitrification and ANAMMOX, nitrification and denitrification, or DIRAMMOX was most frequently identified in both aerobic and anaerobic environments. Notably, among nitrification processes, the partial nitrification converting NH_3_ to NO was more frequently identified as a key reaction than complete ammonia oxidation, despite the Δ_*r*_*G* of partial nitrifications generally being higher than that of complete ammonia oxidation. This prevalence is likely due to the lower oxygen demand per reaction, which allowed faster progression in low oxygen settings, thereby enhancing power generation (Fig. 4c). Meanwhile, partial nitrifications that oxidize NH_3_ to NO_2_^−^ and NO_2_^−^ to NO_3_^−^ contributed significantly to the power only in environments with high oxygen concentrations, as their relatively rapid rates in low O_2_ environments do not offset the disadvantages of their higher Δ_*r*_*G*.

### Microbial nitrogen reaction niches

The variations in the contributions of partial and complete nitrifications to power generation, based on O_2_ level, may partially explain microbial niche specialization. Findings suggest that the relatively higher power achieved by complete ammonia oxidation in low O_2_ conditions supports COMMAMOX *Nitrospira*’s preference for lower O_2_ conditions compared to partial nitrifiers (35, 36). However, at high O_2_ levels, the significant power from the partial nitrification converting NO_2_^−^ to NO_3_^−^ contrasts with reports of the inhibitory effect on the oxidation of NO_2_^−^ by certain NOBs under similar conditions (37). Partial and complete nitrifications utilizing NH_3_ are identified as key reactions even in low pH environments, although the NH_4_^+^/NH_3_ ratio increases as pH decreases. This may partially explain the prevalence of nitrifiers even under acidic conditions (pH < 5.5) (38), while urea spread as fertilizers might serve as an NH_3_ source in low pH fields (39). It has been reported that ANAMMOX in marine environments can occur at O_2_ concentrations higher than the confirmed experimental O_2_ upper limit (approximately 20 µM) (40). The higher power achievement by the combination of ANAMMOX and partial nitrification (reaction 3 in Fig. 2b) suggests that ANAMMOX in such environments can be facilitated by the existence of an O_2_ sink due to the partial nitrification.

Among the key reactions, H^+^ + NH_3_ → NH_4_^+^ and H_2_O + N_2_ + 2.5O_2_ → 2H^+^ + 2NO_3_^−^ are excluded from the community-level network. The latter has the potential to generate proton motive force through the electron transport chain and thus to serve as energy source for ATP synthesis. Since H_2_O + N_2_ + 2.5O_2_ → 2H^+^ + 2NO_3_^−^ contributes significantly to the power only in high oxygen environments (Fig. 4d), if microbial enzymes catalyzing this reaction are discovered in the future, they are likely to be found in aerobic environments.

### Partial reaction utilization: advantages and diversity insurance

Utilizing partial reactions, despite their higher Δ_*r*_*G*, can be energetically beneficial, exemplified by our model prediction. Similar to partial nitrifications, many denitrifiers rely on partial denitrification for ATP synthesis (3, 41). Theoretical studies have shown that under specific conditions, partial reactions proceed more rapidly, thus enhancing ATP production (42–44). Moreover, integrating these reactions within microbial communities boosts overall energetic efficiency and power generation, surpassing the outcome of complete reactions alone (45). The utilization of partial reaction niches may also mitigate the competition for nitrogen sources by allowing them to differentiate niches. This diversity serves as an insurance effect (46), ensuring the resilience and continuity of nitrogen transformation pathways within the nitrogen network.

## Conclusions

Our study provides a comprehensive evaluation of energetically favorable nitrogen reactions and their correspondence with community-level microbial catabolic reactions. Our current nitrogen reaction model includes 988 reactions from 37 nitrogen species half-reactions and O_2_ + 4H^+^ + 4e^−^ ⇌ 2H_2_O. Energetically superior reactions identified by our approach aligned with ANAMMOX and complete and partial nitrification, microbial reactions that rely solely on nitrogen species and oxygen. These results show that the thermodynamic stability of nitrogen species and oxygen significantly explains the microbial community-level catabolic network. By establishing a method for addressing the thermodynamic effect on microbial community metabolism from various redox reactions with minimal chemical parameter setup, our approach can be applied to microbial metabolisms relying on other redox sensitive elements, such as carbon and sulfur. While the simplicity of our model limits its capacity for precise forecasting, the observed alignment between microbial catabolic activities and predictions emphasizes the essential role of thermodynamics in shaping metabolic networks at higher levels of biological organization, providing insight into the universal principles governing biology.

## Methods

### Nitrogen species dynamics model

This model simulates the dynamics of 11 nitrogen species within an open homogeneous single-box aquatic system. The changes in molar concentrations of nitrogen species within the system are influenced by 988 reactions, which are detailed in the subsequent section. The variable vector for the 11 nitrogen species is denoted as **X**_N_ = (NO_3_^−^, NO_2_^−^, NO, N_2_O, N_2_, NH_2_OH, NH_3_OH^+^, N_2_H_4_, N_2_H_5_^+^, NH_4_^+^, NH_3_)^T^, and their molar concentration vector is represented as **M**(t). The dynamics of **M**(t) are governed by the following system of differential equations:

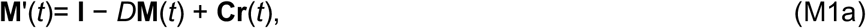

where **I** represents the influx rate vector, *D* the efflux rate constant, **C** the stoichiometric coefficient matrix, and **r**(*t*) the reaction rate vector:

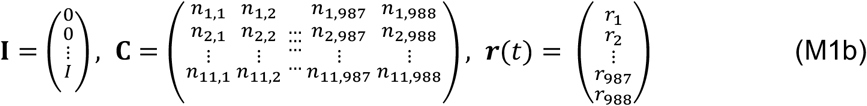

*n*_*k,i*_ represents the stoichiometric coefficient of the *k*th nitrogen species in *rxn*_*i*_. All nitrogen species are exported from the system into the external environment in proportion to their molar concentrations. The concentrations of dissolved oxygen and hydrogen ion were assumed to be constant. The stoichiometric coefficients and reaction rates *r*_*i*_ for each reaction will be detailed in the following sections.

### Reactions involving nitrogen species

The procedure for exhaustively generating all combinations of redox reactions from a group of half-reactions, utilizing the algorithm by (45) as a foundation, was developed in the Wolfram Language (see Supplementary Information and Data). This algorithm was applied to 38 half-reactions, resulting in 962 redox reactions. Reactions were considered distinct if they had different electron transfers, even if their resulting equations were identical. Subsequently, 26 reactions, including acid-base reactions, were added, totaling 988 reactions. For each *rxn*_*i*_ (*i* = 1, …, 988), the stoichiometric coefficient vector is defined as **N**_*i*_ = (*n*_1,*i*_, …, *n*_15,*i*_). *rxn*_*i*_ can then be represented by **N**_*i*_ ·**X**, where **X** is a variable vector that includes **X**_**N**_ along with O_2_, H^+^, and H_2_O. The coefficient matrix **C** in Eq. (M1b) was constructed exclusively from the stoichiometric coefficients for nitrogen species (*n*_1,*i*_, …, *n*_11,*i*_) within **N**_*i*_,, which are transposed and sequenced from the first to the 988th column, forming an 11 × 988 matrix.

### Thermodynamic weight on reaction rates

The reaction rate *r*_*i*_ follows the law of mass action:

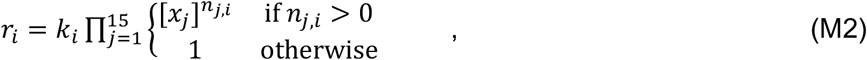

where *k*_*i*_ represents the reaction rate constant for *rxn*_*i*_, [*x*_*j*_] denotes the molar concentration of the *j*th chemical species in the variable vector **X**, and *n*_*j,i*_ is the stoichiometric number of the *j*th chemical species in the *i*th stoichiometric vector **N**_*i*_. If 1 NO_2_^−^ + 0.5 O_2_ → 1 NO_3_^−^ represents *reac*_1_, its reaction rate is given by *r*_1_ = *k*_1_ [NO_2_^−^] [O_2_]^0.5^.

A reaction with Δ_*r*_*G* < 0 proceed relatively quickly compared to its backward reaction and establishes *K*_*eq*_ = exp(-Δ_*r*_*G*º*/*RT) at equilibrium. Based on this relationship, the rate constant *k*_*i*_ for reaction *i* is defined by the following equation:

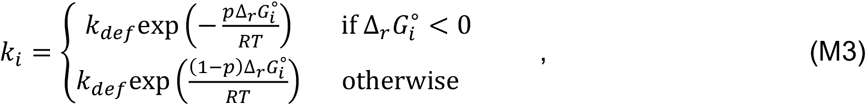

where *k*_*def*_ represents the reference reaction rate constant, and *p* (0 ≤ *p* ≤ 1) is a parameter controlling the thermodynamic weighting on the reaction rate. As *p* approaches 1, the reaction rate constants *k*_*i*_ for reactions with 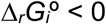 increase exponentially with smaller 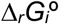 values, while *k*_*i*_ for reactions with 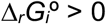 converge to 1. Conversely, as *p* approaches 0, *k*_*i*_ for reactions with 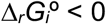 converge to 1, and *k*_*i*_ for reactions with 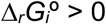 converge to 0 as 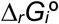 increases.

### Numerical simulation

The set of differential equations in Eq. (M1b) was solved numerically until steady state reached using the NDSolve function in Mathematica 12. The efflux rate constant *D* and the absolute temperature *T* were fixed at 10^−3^ and 288.15K, respectively. Initial values for all nitrogen species were set to 10^−5^ mol L^−1^. Oxygen concentration conditions ranged from −12 ≤ log_10_ [O_2_] ≤ −3 in increments of 0.1, and pH conditions from 2 to 12 in increments of 0.5, totaling 966 conditions for the numerical differentiation calculation. The upper limit of oxygen concentration was set based on the saturation concentration of dissolved oxygen in Earth’s surface waters (≈ 10 mg L^−1^ = 3.1 × 10^−4^ M). Calculations were performed under several conditions for the thermodynamic weighting coefficient *p*, the inflow rate of ammonia *I*, and the reference reaction rate constant *k*_*def*_, to examine the sensitivity of the system to each variable.

## Acknowledgments

This study was supported by JSPS KAKENHI Grant Numbers JP22K06390, JP23H04652, and JP24H01514 to MS, JSPS KAKENHI Grant Number JP22H05153 and JST FOREST program (JPMJFR213E) to OS. JSPS KAKENHI Grant Number 22H05153 to RN. The authors are grateful to the comments from Shawn E. McGlynn and D. Eric Smith.

