## Supplementary Figures for "Thermodynamics Underpinning the Microbial Community-Level Nitrogen Networks"

Supplementary Figure 1

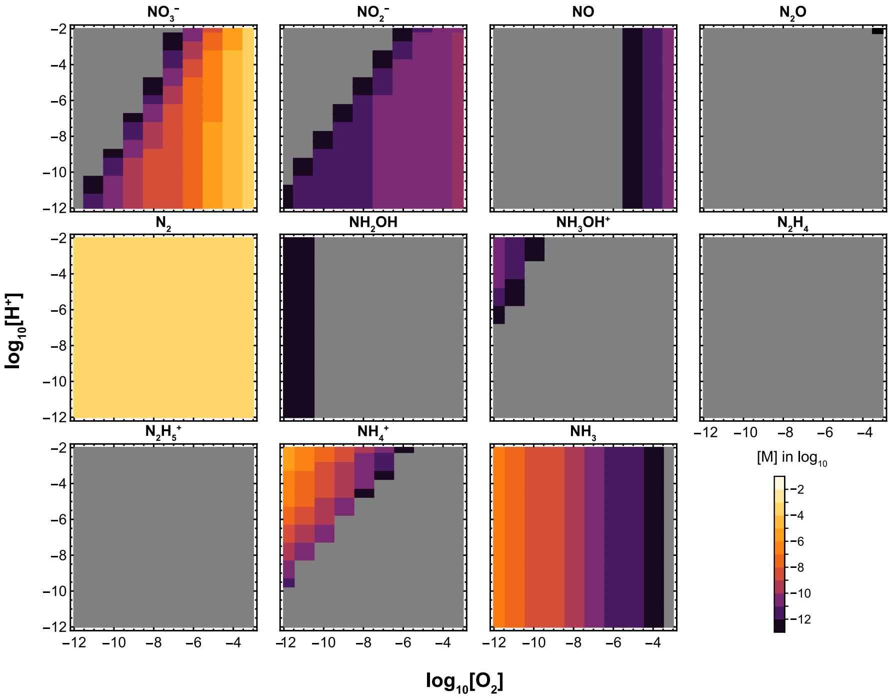


Figure 1: Steady-state concentrations of nitrogens species at various oxygen concentrations and pH levels. The gray areas indicate concentrations below $10^{-12}$ M. $p = 0.2$, $k_{def} = 10^0$, $I = 10^{-5}$, $D = 10^{-3}$, and $T = 288$K

Supplementary Figure 2

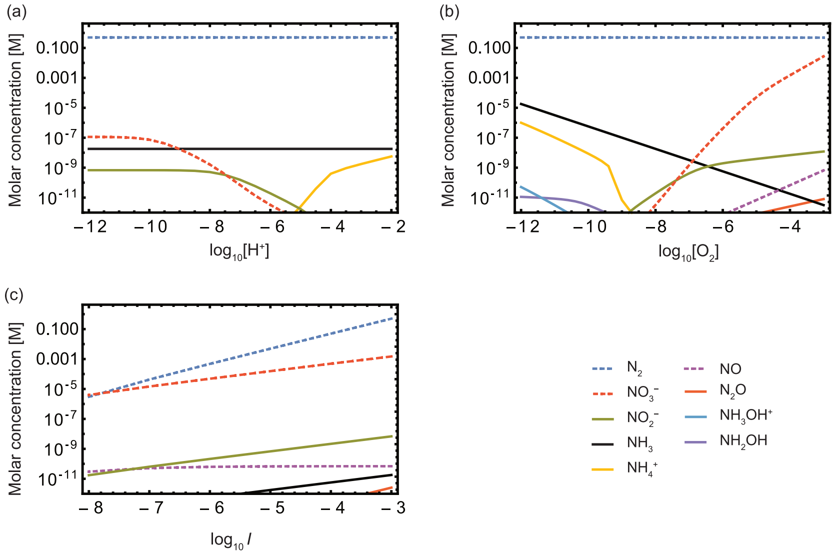


Figure 2: Steady state response of nitrogen species concentrations to variations in hydrogen ion concentration (a), oxygen concentration (b), and ammonia influx (c). (a) $[O_2] = 10^{-8}$ and $I = 10^{-3}$, (b) $[H^+] = 10^{-6}$ and $I = 10^{-3}$, (c) $[O_2] = 10^{-4}$ and $[H^+] = 10^{-6}$. Other parameter values are $p = 0.2$, $k_{\text{def}} = 10^0$, $D = 10^{-3}$, and $T = 288$.

Supplementary Figure 3

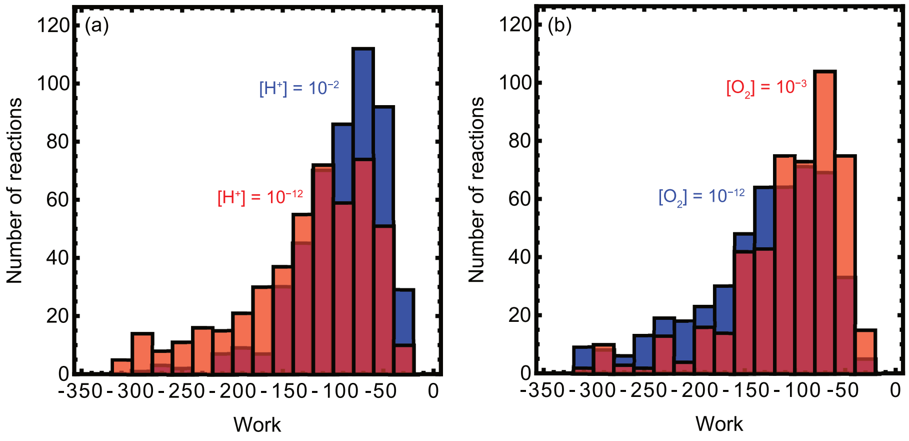


Figure 3: Distribution of the steady-state power generation $P_i = -\Delta_r G_i \times r_i$ for 988 reactions. (a) $[O_2] = 10^{-3}$ M, (b) pH = 8. Other parameter values are $I = 10^{-3}$, $p = 0.2$, $k_{\mathrm{def}} = 10^{-10}$, $D = 10^{-3}$, and $T = 288$ K.
